# Genome editing for treatment of *JAK2* V617F-driven myeloproliferative neoplasms

**DOI:** 10.64898/2025.12.17.694595

**Authors:** Roman M. Doll, Yuqi Shen, Zoe C. Wong, Ramy Slama, Maria Greco, Lai Cheng, Eleni Louka, Weijiao Zhang, Yavor Bozhilov, Joseph C. Hamley, Adam C. Wilkinson, Bethan Psaila, Adam J. Mead, James O.J. Davies

## Abstract

*JAK2* V617F is a common haematological driver mutation and underlies most cases of myeloproliferative neoplasms (MPNs). Reducing variant allele frequency (VAF) is an important treatment goal, but no current treatment modalities fully and specifically inhibit mutant signalling. Thus, the consequences of V617F inactivation are unclear, including whether selective inhibition of V617F signalling will eradicate mutant cells due to oncogene addiction. Here, we describe an allele-selective CRISPR-Cas genome editing strategy that achieves selective and efficient *JAK2* V617F inactivation in patient stem and progenitor cells. We show that *JAK2* V617F heterozygous cells are not oncogene addicted and maintain viability and differentiation potential upon loss of their mutant allele. In contrast, homozygous mutant cells are eradicated upon deletion of both mutant alleles. Across *in vitro*, organoid, xenotransplantation and single-cell assays, selective deletion of V617F alleles reverts MPN hallmarks including erythroid clonogenicity, inflammatory gene expression signatures, as well as myelofibrosis and splenomegaly phenotypes in an *in vivo* xenograft model. Collectively, our results show that is it possible to revert heterozygous *JAK2* V617F mutant cells to a normal phenotype and suggest that *ex vivo* genome editing of stem and progenitor cells may be a viable treatment option to achieve rapid and deep reductions in VAF.

## Introduction

Myeloproliferative neoplasms (MPNs) are stem cell-propagated blood cancers that are driven in most cases by the *JAK2* V617F mutation, often as the sole genetic event.^1^ Molecular response, measured as a reduction in *JAK2* V617F variant allele frequency (VAF), is emerging as an important treatment goal and major predictor of favourable long-term outcomes.^2^ However, current JAK2 inhibitors do not provide complete inhibition of mutant signalling and do not eliminate the mutant clone in most patients.^2,3^ Moreover, since patients frequently carry a mix of both heterozygous and homozygous mutant cells,^4^ it is unknown how hematopoietic stem and progenitor cells (HSPCs) of different genotypes would respond to ablation of mutant signalling. Additionally, given that niche effects are major drivers of MPN,^5,6^ it is unclear whether inhibition of V617F alone is sufficient to revert disease hallmarks. Although a mouse model with a reversible V617F mutation has previously been described,^7^ it has thus far not been possible to address these questions in human patient cells.

Here, we developed specific and efficient allele-selective genome editing of *JAK2* V617F which allowed us to investigate the effects of mutant signalling inhibition in primary CD34+ cells. Across *in vitro*, organoid, xenotransplantation and single-cell assays, we demonstrate that heterozygous mutant cells are not oncogene addicted and that genetic inactivation of V617F alleles is sufficient to revert MPN hallmarks. Given the remarkable efficiency and specificity of our targeting strategy and the decade long latency that is required for a low number of mutant cells to initiate disease,^8^ our results furthermore suggest that *ex vivo* genome editing in an autologous transplant setting may be a viable treatment option for V617F-driven MPNs.

## Results

### Highly efficient and specific JAK2 V617F inactivation using Cas12a

We reasoned that allele-selective genome editing of V617F alleles would counteract mutant gene effects while preserving normal haematopoiesis from wild-type *JAK2* copies.^9^ We developed a strategy based on a Cas12a target site which uses a *de novo* protospacer adjacent motif (PAM) created by the V617F mutation.^10^ Crucially, this gRNA places the nuclease cut site within the adjacent conserved splice donor motif (Supplementary Fig. 1A), suggesting that CRISPR indels will selectively inactivate mutant gene copies by interfering with splicing. We electroporated Cas12a Ultra ribonucleoproteins (RNPs) into SET-2 cells, which harbour V617F alleles at a VAF of ~70%, and used amplicon sequencing to sensitively characterize the editing outcomes at the V617F locus. Cas12a Ultra-targeting resulted in an average of 96.6% indels on mutant gene copies (Supplementary Fig. 1B), substantially outperforming a previously described Cas9-based allele-selective gRNA both in terms of editing efficiency and favourability of the indel profile^11,12^ (Supplementary Fig. 1C,D). Moreover, allele-selective Cas12a Ultra-targeting of V617F abrogated the cytokine-independent growth of SET-2 cells (Supplementary Fig. 1E,F) and no alleles other than unedited ones enriched over time (Supplementary Fig. 1G), further confirming that Cas12a targeting functionally inactivates V617F copies.

We also performed GUIDE-Seq^13^ to determine the off-target profile of this editing strategy. In line with the previously reported high genome-wide specificity of Cas12a nucleases, we only observed two off-targets with low read counts (Supplementary Fig. 1H), both of which reside within non-coding regions without features of regulatory chromatin (Supplementary Fig. 1I,J).

### JAK2 V617F inactivation in primary CD34+ cells dramatically reduces VAF but preserves wild-type and heterozygous cells

We then employed our Cas12a Ultra editing strategy on CD34+ cells from polycythaemia vera (PV) or post-PV myelofibrosis (PPV-MF) patients with single-hit V617F mutation. As the number of cells obtained from a typical sample was generally an order of magnitude below those required for electroporation, we leveraged a recently described polymer-based expansion culture to expand the CD34+ cells prior to electroporation^14^ (Supplementary Fig. 1K). Editing across four donors was highly robust and selective with an average efficiency of 96.2% on mutant and 2.7% on wild-type alleles (Fig. 1A), leading to a drastic drop in VAF after only 96 hours (Fig. 1B).

**Fig. 1:**
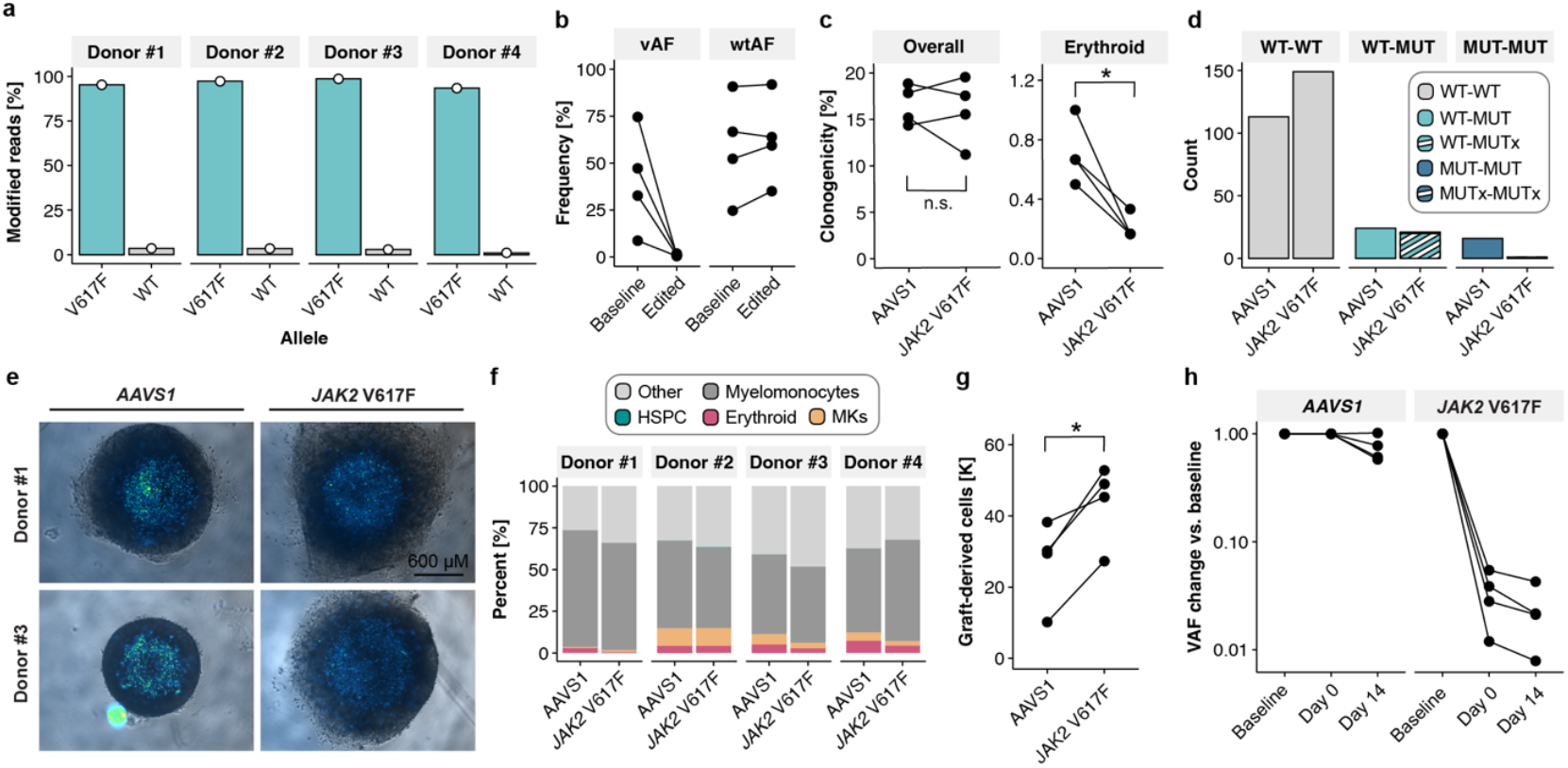
Allele-selective V617F inactivation in primary CD34+ cells via genome editing. **a**, Editing efficiencies of WT and V617F alleles 96h after electroporation of patient CD34+ cells in n = 4 independent patient samples, determined via amplicon sequencing. **b**, Frequency of unedited V617F alleles (vAF) and wild-type alleles (wtAF) in *AAVS1* (baseline) or *JAK2* V617F (edited) edited cells from n = 4 independent patient samples. **c**, Percentage of CD34+ cells forming any colony, or erythroid colonies in n = 4 independent patient samples. p-values calculated via paired two-sided t-test, n.s.: p > 0.05, *: p <= 0.05. **d**, Number of colonies with indicated genotype, with x indicating a mutant allele that has been disrupted by a CRISPR indel, total from n = 4 independent patient samples. **e**, Representative images of bone marrow organoids three days post engraftment with labelled CD34+ cells (in total n = 4 edited patient samples were engrafted). Organoid shown as brightfield and Celltrace Violet labelled patient-derived cells in firefly scale. Images were taken on an EVOS M5000 system. **f**, Flow cytometry analysis of patient-derived cells 14 days post organoid engraftment (n = 4 independent patient samples). **g**, Number of graft-derived cells at 14 days post engraftment into bone marrow organoids. p-values calculated via paired two-sided t-test, *: p <= 0.05. **h**, VAF (i.e. frequency of unedited V617F amplicon sequencing reads) at baseline (*AAVS1* edited day 0) versus day 0 and 14 of organoid engraftment in n = 4 independent patient samples.

Following editing of *JAK2* V617F, HSPCs retained colony forming potential with the expected reduced erythroid output compared to control edited cells (*AAVS1*, Fig. 1C). Given the essentiality of *JAK2* and the fact that patients frequently carry a mix of heterozygous and homozygous mutant cells owing to loss of heterozygosity events,^4^ we wanted to assess whether inactivation of V617F is differentially tolerated depending on mutant zygosity. To this end, we genotyped 342 individual colonies and found that while heterozygous mutant cells continued to form colonies with an inactivated copy of their V617F allele, homozygous mutant cells did not (Fig. 1D).

Next, we transplanted edited and fluorescently labelled CD34+ cells into bone marrow organoids^15^ which showed normal engraftment (Fig. 1E) and multilineage differentiation of myeloid cells (Fig. 1F). Overall cellular output was increased compared to control edited cells (Fig. 1G), potentially due to counteracting detrimental cell extrinsic effects of mutant cells on the organoid niche. Crucially, VAF was substantially reduced by editing and reached as little as 1-2% of the baseline allele burden at the experimental endpoint (Fig. 1H). These findings support that HSPCs with at least one wild-type *JAK2* copy retain colony forming and differentiation potential after selective editing of V617F mutant alleles, and that deep reductions in VAF can persist through differentiation and extensive proliferation.

### Combined single cell genotyping and RNA-Seq shows reversal of JAK2 V617F-driven transcriptional signatures

Next, we performed single cell combined allelic-level genotyping and RNA sequencing of edited CD34+ cells (TARGET-Seq) to determine the impact of allele-specific *JAK2* V617F genome editing on wild-type versus *JAK2* V617F heterozygous or homozygous HSPCs (Fig. 2A).^16,17^ We obtained high-quality genotyping and transcriptome data from 1175 cells across four donors (Supplementary Fig. 2A-C) and identified the initial genotype of wild-type (WT-WT), heterozygous (WT-MUT) and homozygous mutant (MUT-MUT) cells (Fig. 2B). Projection onto a human bone marrow reference^18^ revealed a diverse set of hematopoietic progenitor cells, with a notable population of EoBasoMast precursors (EBMs, Fig. 2C), as recently described.^6^ *JAK2* VAF was lowest in HSCs and increased towards lineage-committed progenitors (Supplementary Fig. 2D). Investigating *JAK2* V617F editing outcomes in single cells showed minimal occurrence of undesired genotypes such as mono-allelically edited homozygous mutant cells (MUT-MUTx, with x referring to an allele carrying a CRISPR indel) or edited wild-type gene copies (WT-WTx, Supplementary Fig. 2E). Overall, we found that 97.2% (377 out of 388) of mutant cells subjected to V617F-targeting RNPs had all mutant alleles inactivated (Fig. 2D).

**Fig. 2:**
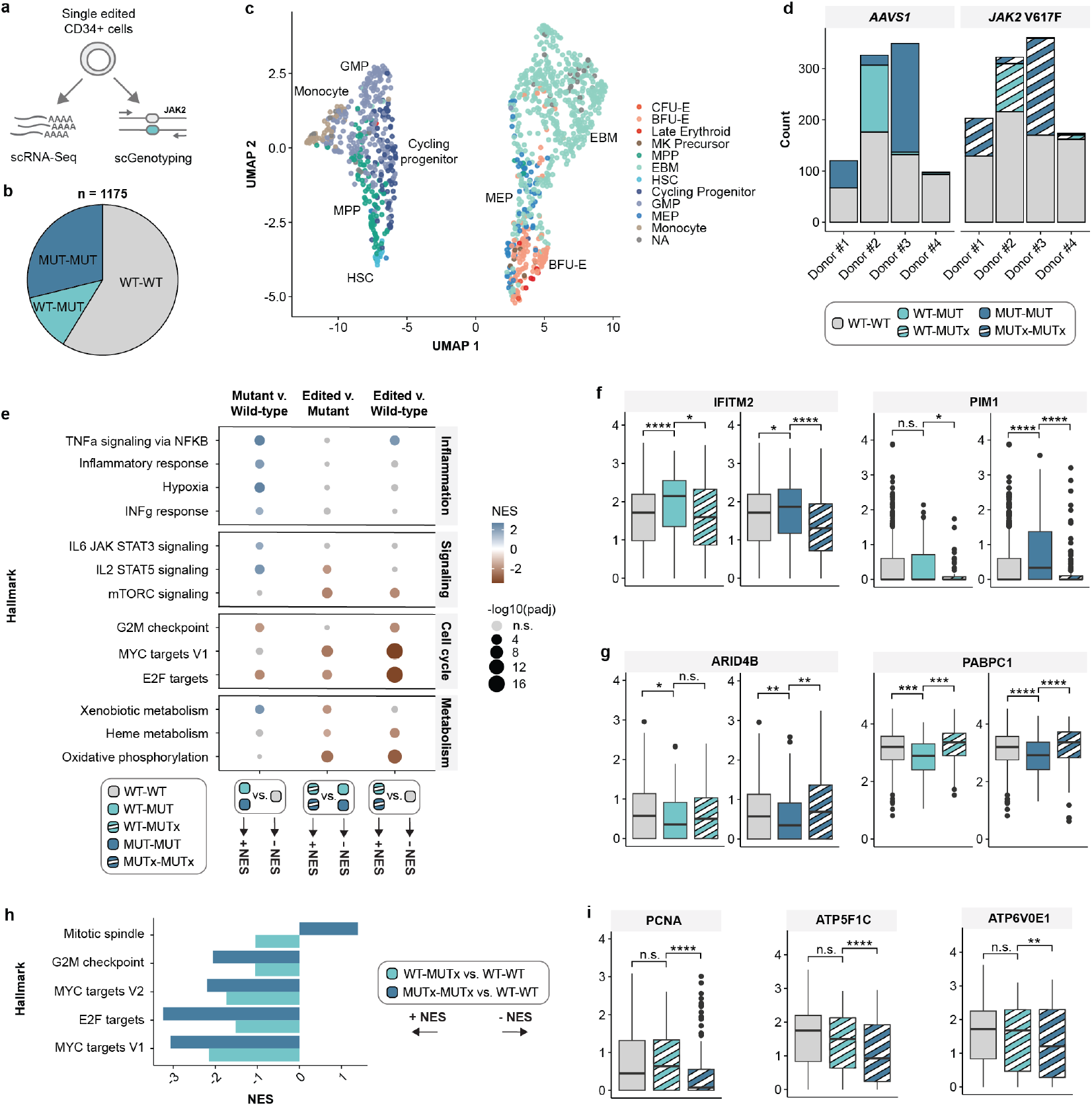
Transcriptional consequences of V617F inactivation on HSPCs at single cell resolution. **a**, Schematic representation of TARGET-Seq. **b**, Number of cells with corresponding genotypes that passed all quality filtering steps. **c**, UMAP of cell types inferred from projection onto an adult bone marrow reference dataset. **d**, Number of cells with indicated genotypes across donors and editing conditions. **e**, GSEA for selected hallmarks grouped by WT (i.e. WT-WT), MUT (i.e. both WT-MUT and MUT-MUT) and MUTx (i.e. both WT-MUTx and MUTx-MUTx) cells. **f**, Log-normalized expression of direct JAK-STAT target genes. **g**, Log-normalized expression of regulators of erythroid differentiation. **h**, Comparison of NES scores for cell-cycle associated pathways between heterozygous and homozygous conditions. **i**, Log-normalized expression of PCNA and ATP Synthetase subunits. For all gene expression plots, p-values calculated via two-sided Wilcoxon test. n.s.: p > 0.05, *: p <= 0.05, **: p <= 0.01, ***: p <= 0.001, ****: p < = 0.0001. All data obtained from a total of n = 4 independent patient samples.

Next, we investigated the effects of the V617F mutation and its inactivation on gene expression. Differential expression (Supplementary Fig. 2F-G) and gene set enrichment analysis (GSEA, Fig. 2E and Supplementary Table 1) between WT-WT and any mutant allele-bearing cell (WT-MUT and MUT-MUT) revealed that mutant cells were enriched in inflammatory and signalling pathway signatures, which was largely reverted upon inactivation of V617F. Importantly, V617F inactivation reduced the expression of JAK-STAT target genes such as *PIM1* and *IFITM2* in both heterozygous and homozygous mutant cells (Fig. 2F). Furthermore, following editing, mutant cells were depleted in cell cycle-associated expression signatures including E2F targets. Mutant cells also exhibited altered expression of *PABPC1* and *ARID4B*, which are essential for normal erythroid differentiation (Fig. 2G).^19,20^ Crucially, V617F inactivation reverted expression of both *PABPC1* and *ARID4B* to the levels found in wild-type cells (Fig. 2G), suggesting that inactivation of the V617F allele is sufficient to mitigate aberrant *JAK2* V617F-associated gene expression defects in erythropoiesis *ex vivo*.

Next, we sought to uncover differential effects of V617F inactivation on heterozygous versus homozygous mutant cells. GSEA split by WT-MUT or MUT-MUT genotype showed that while V617F inactivation showed marked depletion of cell-cycle signatures in homozygous cells, this effect was substantially less pronounced in heterozygous mutant cells (Fig. 2H, Supplementary Fig. 2H and Supplementary Table 2). Indeed, while MUTx-MUTx cells exhibited lower expression of the proliferation marker *PCNA* and several ATP synthetase subunits, WT-MUTx maintained expression levels comparable to WT-WT cells (Fig. 2I), in keeping with our observation in functional experiments that WT-MUTx but not MUTx-MUTx cells maintain viability post editing.

### JAK2 V617F inactivation exerts therapeutic benefit in a xenograft model of MPN

Finally, we set out to investigate the effects of *JAK2* V617F inactivation in an *in vivo* model. We electroporated SET-2 cells with RNPs targeting either *JAK2* V617F (henceforth ‘treatment’) or *AAVS1* (henceforth ‘control’) and transplanted the cells into sub-lethally irradiated NSG mice (Fig. 3A). While control animals rapidly developed leukemia, survival was significantly improved in the treatment arm (Fig. 3B). At weeks 6-8, we observed pronounced splenomegaly in control animals which was fully reverted upon *JAK2* V617F inactivation (Fig. 3C,D). In control mice, immunohistochemistry revealed a fibrotic bone marrow with limited cellular diversity and complete absence of megakaryocytes (Fig. 3E). Conversely, marrow of treatment group animals appeared histologically normal and did not exhibit any signs of reticulin fibrosis. Flow cytometry detected human cells in both spleen and bone marrow of treatment group animals but significantly below those levels observed in the control group (Fig. 3F). Importantly, the SET-2 cells that were present in the bone marrow and spleen of treatment animals exhibited *JAK2* VAFs of less than 1% (Fig. 3G). Collectively, these findings show that allele-selective *JAK2* V617F inactivation reverts MPN phenotypes *in vivo* while preserving viability and engraftment potential of cells harbouring a wild-type copy of JAK2.

**Fig. 3:**
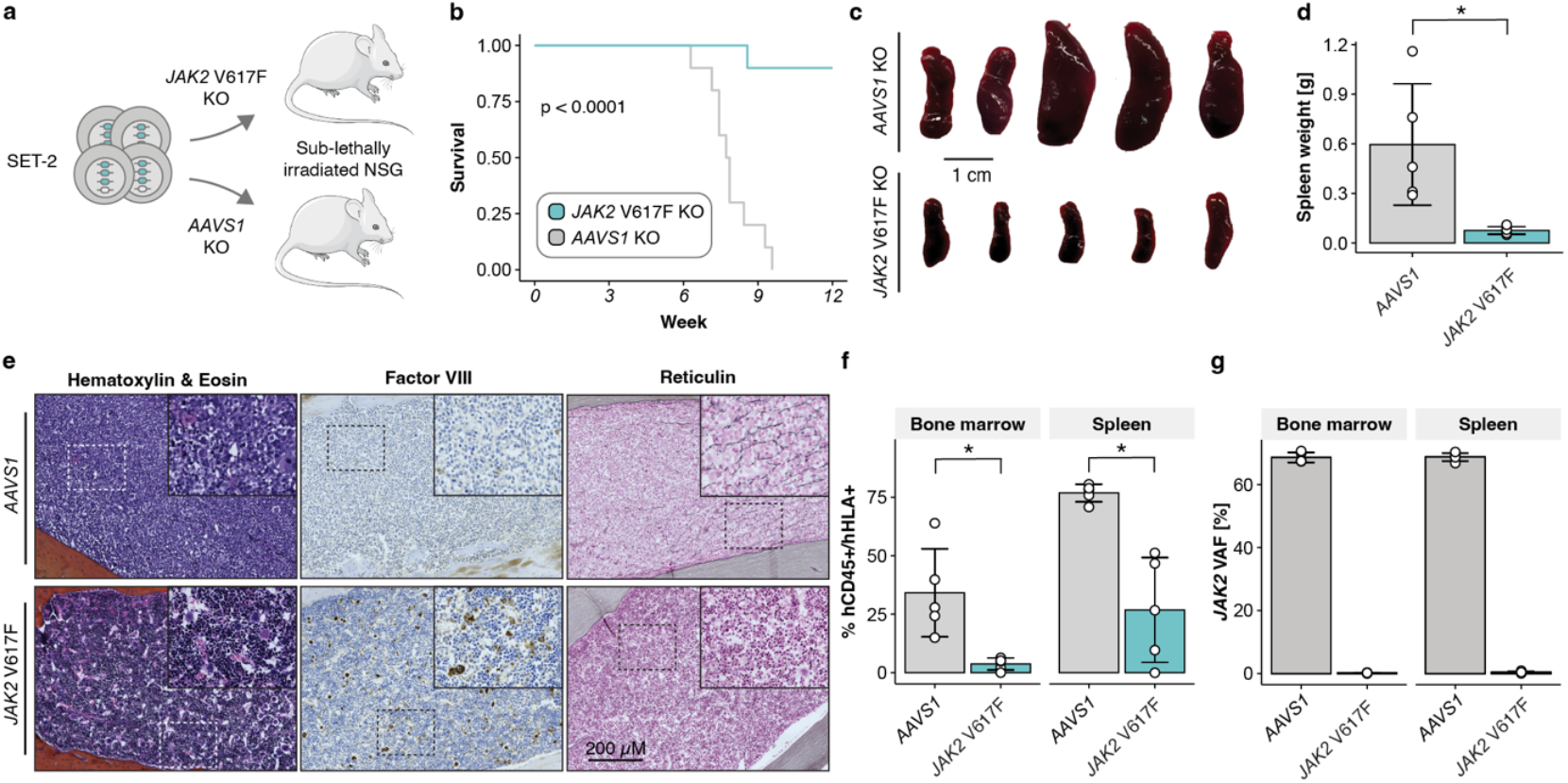
Allele-selective V617F inactivation exerts therapeutic benefit in a xenograft model of MPN. **a**, Schematic of xenograft experiment. **b**, Survival curves from N=10 animals per group from two independent transplant cohorts. p-value calculated via log rank test. **c**, Spleen images. **d**, Spleen weights corresponding to c (n = 5 per group, mean±s.d). p-values calculated via two-sided paired t-test, n.s.: p > 0.05, *: p <= 0.05. **e**, Representative images of bone marrow histology for general bone marrow morphology (H&E), megakaryocytes (Factor VIII) and fibrosis (reticulin). Images were acquired on a Zeiss Axioscan 7 at 10x magnification. **f**, Flow cytometry analysis of human chimerism (n = 5 per group, mean±s.d). p-values calculated via two-sided paired t-test, n.s.: p > 0.05, *: p <= 0.05. **g**, *JAK2* VAF, i.e. percent human amplicon sequencing reads with unedited V617F alleles (n = 5 per group, mean±s.d). Unless otherwise indicated, data from animals culled between weeks 6-8 after transplantation.

## Discussion

We have established highly efficient allele-selective editing of *JAK2* V617F in primary patient CD34+ cells, allowing us to elucidate the distinct effects of ablation of mutant signalling in heterozygous and homozygous mutant cells. We show that the heterozygous V617F mutation does not induce oncogene addiction^21^ and that heterozygous cells can thus be reverted to a normal phenotype. Our findings contrast with a recently described mouse model where deletion of a heterozygous V617F mutation caused dramatic reductions in mutant cell fraction,^7^ suggesting that murine models may not faithfully recapitulate human disease biology. Importantly, our observations predict that JAK2 mutant-selective inhibitors, which are currently in clinical development, will fail to eradicate cells harbouring heterozygous JAK2 mutations as these cells will revert to a normal hematopoietic phenotype. Nevertheless, since mutant allele inactivation was sufficient to counteract MPN hallmarks both *in vitro* and *in vivo*, our findings suggest that selective ablation of *JAK2* V617F is sufficient to reverse disease phenotypes even in the setting of advanced disease such as V617F-driven myelofibrosis.

Beyond generating insights into V617F biology, our study also provides a potential novel therapeutic avenue. Unlike current JAK2 inhibitors, our genome editing approach achieves rapid and deep reductions in VAF, which is emerging as an important treatment goal.^2^ Given the increasing adoption of *ex vivo* CRISPR editing of CD34+ cells in an autologous transplant setting for mendelian diseases,^22^ we propose that a similar approach may be developed for the treatment of V617F-driven MPNs. Although *ex vivo* editing rates of 100% are currently unattainable, the ~97% reduction in clonal burden achieved by our strategy is likely to result in substantial long-term benefit by resetting clonal expansion by several decades^8^ and shrinking the pool of cells that can transform to more severe malignancy.

## Methods

### Human tissue

Peripheral blood and bone marrow samples were collected from MPN patients under the approval of the INForMeD Study (REC: 199833, 26 July 2016, University of Oxford). Patients and normal donors provided written informed consent in accordance with the Declaration of Helsinki for sample collection and use in research. Cells were subjected to Ficoll gradient centrifugation and for some samples, CD34 enrichment was performed using immunomagnetic beads (Miltenyi). Total mononuclear cells (MNCs) or CD34+ cells were frozen in FBS supplemented with 10% DMSO for further analysis. Cryopreserved peripheral blood mononuclear cells stored in FCS with 10% DMSO were thawed and processed by warming briefly at 37°C, gradual dilution into IMDM supplemented with 10% FCS and 0.1 mg/mL DNase I, centrifuged at 500 g for 5 minutes and washed in FACS buffer (PBS + 2mM EDTA + 10%FCS).

### Cell culture and electroporation

SET-2 cells were cultured in RPMI-1640 supplemented with 20% FBS and 1% Penicillin-Streptomycin. Patient CD34+ cells were either thawed directly (see above) or selected from thawed MNC samples using the CD34 MicroBead Kit UltraPure (Miltenyi, see Supplementary Table 3 for a list of patient samples used in this study). The CD34+ cells were then expanded in a polymer-based expansion system for two weeks as previously described,^14^ with the addition of 10 ng/µL recombinant stem cell factor (SCF). Electroporation was performed using the 4D nucleofector (Lonza) according to manufacturer’s guidelines, using 1.7 µL Cas9 or Cas12a nuclease (IDT), 160 pmol crRNA (IDT) for Cas12a and 160 pmol sgRNA (Synthego) for Cas9, as well as the corresponding electroporation enhancers (IDT). gRNA sequences are listed in Supplementary Table 4. SET-2 cells were electroporated using the SF kit and pulse code FF-120. Primary CD34+ cells were electroporated using the P3 kit and pulse code DS-130 and then recovered for 48 hours in StemSpan SFEM II supplemented with 100 ng/µL of recombinant SCF, TPO and FLT3L, followed by another 48 hours recovery with supplementation of 25 ng/µL SCF and 1 ng/µL TPO before downstream assays were commenced.

### Colony-forming assays

Ten live CD34+ cells were sorted into each well of a 96-well plate containing 50 µL Methocult media (Stem Cell Technologies), using one plate of H4435 and H4531 for each sample and editing condition. Colonies were scored 12-14 days after sorting. Erythroid clonogenicity was determined from EPO-containing plates only (H4435) as we did not detect EPO-independent colony formation in any of the samples using H4531. Individual colonies were then picked into PBS, centrifuged at 500 g for 5 minutes, and resuspended in 10 µL quick lysis buffer (10 mM Tris-HCl pH 7.5, 0.05% SDS and 0.3% Proteinase K, NEB, molecular biology grade). The samples were then incubated at 65°C for 5 minutes, followed by heat inactivation at 98°C for 2 minutes. The resulting whole cell lysate was used as input for amplicon sequencing of the *JAK2* locus, using three rounds of PCR (first, preamplification of a 500 bp target region, second, a nested PCR adding partial Illumina TrueSeq sequences and plate barcodes and third, generation of Illumina-ready amplicons by amplification of the product with NEBNext 96 unique dual index set, E6440 or E6442). Primer sequences and PCR conditions are described in Supplementary Table 5.

### Bone marrow organoids

Routine iPSC cultures were maintained on Geltrex (ThermoFisher Scientific)-coated plates in StemFlex (ThermoFisher Scientific) media with clump passaging performed using ReLeSR (Stem Cell Technologies). Cells were tested for mycoplasma at 3-month intervals, with karyotyping performed at least every 12 months. Bone marrow organoids were generated as previously described^15,23^ using a fluorescent iPSC line, MCND-TENS2-mScarlet3 (parental line registered at https://hpscreg.eu/cell-line/RTIBDi001-A). On day 14 of organoid differentiation, primary cells were labelled with CellTrace Violet (Invitrogen) following kit instructions. Labelled primary cells were seeded into organoids at a density of 5,000 cells per organoid. Engrafted organoids were cultured in StemPro-34 SFM medium (ThermoFisher Scientific) supplemented with 2% KnockOut serum (ThermoFisher Scientific), 2% Chemically Defined Lipids (ThermoFisher Scientific), 0.5% Pen/Strep, and cytokines (5 ng/mL of EPO and 1 ng/mL of TPO). Engrafted organoids were imaged using EVOS M5000 system (ThermoFisher Scientific) to monitor CellTrace Violet-labelled cell engraftment within the organoids.

Fourteen days after engraftment, organoids were collected by gravitation and gently washed with PBS to remove non-engrafted cells. Multiple organoids were pooled together by condition and dissociated using 2.5 mg/mL collagenase D (Roche) dissolved in HBSS (Sigma) and supplemented with 1% FBS and 1000 U/mL DNase I in IMDM medium, as previously described.^24^ Organoids were incubated at 37°C for 15-30 minutes and gentle trituration with a p200 pipet was performed every 6-8 minutes. Once a single-cell suspension was obtained, the dissociation was stopped using 10% FBS, 20 mM EDTA in PBS. Cells were resuspended, stained with the following antibodies: anti-hCD45-APC-eF780 (BioLegend, Cat# 368516), anti-hCD34-BV650 (BioLegend, Cat# 343624), anti-hCD71-BV510 (BD Biosciences, Cat# 744926), anti-hCD235-PE-Cy7 (BioLegend, Cat# 306620), anti-hCD41-FITC (BioLegend, Cat# 303704), anti-hCD42b-AF647 (BioLegend, Cat# 303924), anti-hCD11b-AF700 (BioLegend, Cat# 301355), anti-hCD14-AF700 (BioLegend, Cat# 367113). Samples were analysed on a BD FACSAria Fusion cell sorter, and mScarlet-negative primary cells were sorted for VAF analysis. Flow cytometry data were analysed using FlowJo (v10.10) software.

### Mouse xenotransplantation

Animals were bred and maintained in accordance with UK Home Office regulations and all experiments conducted in accordance with approvals from the University of Oxford Animal Welfare and Ethical Review Body (project license PP3929317). One day post-electroporation, 500,000 SET-2 cells were transplanted via tail vein injection into sub-lethally irradiated (2 Gy) 8–12-week-old NSG males. Animals were culled upon loss of 15% of the initial body weight or when paralysis of the limbs was apparent. Human chimerism of bone marrow and splenocytes was assessed via flow cytometry using antibodies anti-hCD45-BV421 (Biolegend, Cat# 304032), anti-hHLA-ABC-FITC (BD Biosciences, Cat# 555552) and anti-mouse CD45.1-PE/Cy7 (Biolegend, Cat# 110730). *JAK2* VAF was assessed via amplicon sequencing of DNA extracted from bone marrow and splenocytes respectively, as described below. For immunohistochemistry, leg bones were fixed in formaldehyde and subsequently sectioned and stained at Gustave Roussy Cancer Campus, PETRA.

### Amplicon-sequencing and data analysis

For bulk amplicon sequencing, whole cell lysates where prepared by resuspension of the cell sample in quick lysis buffer (10 mM Tris-HCl pH 7.5, 0.05% SDS and 0.3% Proteinase K, NEB, molecular biology grade), followed by incubation at 65°C for 5 minutes and heat inactivation at 98°C for 2 minutes. Sequencing libraries were prepared using a 2-stage PCR protocol comprised of a first PCR amplifying the target region with primers encoding partial Illumina TrueSeq adapters, followed by generation of Illumina-ready amplicons by amplification of the first PCR product with NEBNext 96 unique dual index set, E6440 or E6442. Primer sequences and PCR conditions are described in Supplementary Table 6. Amplicons were sequenced with Illumina 150 bp paired-end reads and the data analysed using CRISPresso2^25^ in allele-selective mode. *JAK2* VAF was calculated as the overall percentage of sequencing reads encoding unedited *JAK2* V617F alleles.

### GUIDE-Seq

GUIDE-Seq was performed in SET-2 cells via electroporation of RNPs and dsODN. Library preparation, sequencing and data analysis was performed as previously described^13^ (see Supplementary Table 7 for sample barcoding and analysis information).

### TARGET-Seq+ library preparation and data analysis

We sorted a total of 2,280 CD34+ cells from four donors edited with either *JAK2* V617F or *AAVS1* targeting RNPs into 384-well plates and prepared amplicon and transcriptome libraries as previously described^16,17^ (see Supplementary Table 8 for a list of primers).

For analysis of the genotyping data, the sequencing reads were demultiplexed and analysed with CRISPResso2^25^ for each individual cell. Cells with fewer than 500 supporting reads were excluded from downstream analysis (cutoff determined based on read counts from blank wells). We first called the pre-editing genotype of each cell based on the percentage of reads aligning to either *JAK2* WT or *JAK2* V617F using a cutoff of 12.5% and 87.5% for heterozygosity and homozygosity, respectively. We then determined editing outcome based on the percentage of wild-type or mutant alleles carrying indels. For heterozygous cells, only cells with more than 80% editing were called as edited. For homozygous cells, given the possibility of only one of the two mutant alleles being edited, we called cells as mono-allelically edited (i.e. MUT-MUTx) if the editing percent was between 15% and 85%, and bi-allelically edited (i.e. MUTx-MUTx) if it was >85%. Cutoffs were determined empirically to avoid misclassifications caused by index hopping or other sources of cross-contamination, which was apparent through cells exhibiting diverse indel products, which is implausible given the fact that each cell can at most harbour two distinct indels at the target site.

Demultiplexing, adapter trimming, alignment and feature counting of the transcriptome sequencing data was performed as previously described.^17^ Using Seurat v5.2.1^26^, the dataset was filtered for cells with at least 20,000 reads, 2000 features and less than 15% mitochondrial RNA reads, followed by standard normalization, variable feature detection and data scaling, including regression of cell cycle scores. For inference of cell types, the data was projected on a recently published single cell bone marrow atlas, using the BoneMarrowMap R package v0.1.0.^18^ Differential expression analysis was performed based on the Wilcoxon rank sum test. Genes were prefiltered for protein coding genes, an overall detection rate in at least twenty percent of cells, and a minimum log-fold change of 0.2. GSEA was performed using fgsea v1.32.2.

## Author contributions

**R.M.D**. Conceptualization, investigation, data analysis, visualization, writing (original draft and editing). **Y.S**.: Investigation, data analysis. **Z.C.W**.: Investigation, data analysis. **R.S**.: Investigation, **M.G**.: Investigation. **L.C**.: Investigation. **E.L**.: Investigation. **W. Z**.: Investigation. **Y.B**.: Investigation. **J.C.H:** Data analysis. **A.C.W:** Methodology. **B.P:** Methodology. **A.J.M**: Conceptualization, resources, supervision, funding acquisition, methodology, writing (original draft and editing), project administration. **J.O.J.D**.: Conceptualization, resources, supervision, funding acquisition, methodology, writing (original draft and editing), project administration. All authors read and approved the submitted manuscript.

## Data availability

TARGET-Seq and GUIDE-Seq data have been deposited in the GEO databank (GSE300279 and GSE300280). Custom code for TARGET-Seq data analysis is available on GitHub (https://github.com/Davies-Genomics-and-Genome-Editing-Lab/Doll_et_al_Scripts)

## Acknowledgments

R.M.D. is supported by the Clarendon Fund, a Balliol College John Henry Jones scholarship, a Scatcherd Scholarship, and Radcliffe Department of Medicine Studentship. J.O.J.D is supported by the MRC Molecular Haematology Unit (MC_UU_00029/04), Oxford NIHR Biomedical Research Centre, NIHR Blood and Transplant Research Unit in Precision Therapeutics, and Wellcome (225220/Z/22/Z). W.Z. was supported by an MRC project grant (MR/T030410/1) and the MRC Molecular Haematology Unit (MC_UU_00029/04). This work was supported by a CRUK Senior Cancer Research Fellowship to A.J.M. (Grant number C42639/A26988) and a CRUK Discovery Programme Award to A.J.M. (Grant number DRCNPG-Nov24/100003). The authors would like to acknowledge the CRUK Oxford Centre (CTRQQR-2021\100002), the National Institute for Health Research (NIHR) Oxford Biomedical Research Centre (BRC); John Fell Fund (131/030 and 101/517), the EPA fund (CF182 and CF170), the MRC WIMM Strategic Alliance awards G0902418 and MC_UU_12025, and the contribution of the WIMM Sequencing Facility, supported by the MRC Human Immunology Unit and by the EPA fund (CF268). We would also like to acknowledge the WIMM Advanced Single Cell OMICS Facility, the WIMM Flow Cytometry Core Facility, the WIMM Genome Engineering Core Facility as well as the Biomedical Services (BMS) of the University of Oxford. The views expressed are those of the authors and not necessarily those of the National Health Service (NHS), the NIH, the NIHR or the Department of Health and Social Care.

## References

1 Mead, A. J. & Mullally, A. Myeloproliferative neoplasm stem cells. Blood 129, 1607–1616 (2017). 10.1182/blood-2016-10-696005

2 Harrison, C. N. et al. Ruxolitinib Versus Best Available Therapy for Polycythemia Vera Intolerant or Resistant to Hydroxycarbamide in a Randomized Trial. J Clin Oncol 41, 3534–3544 (2023). 10.1200/JCO.22.01935

3 Gorantla, S. P. et al. Efficacy of JAK1/2 inhibition in murine myeloproliferative neoplasms is not mediated by targeting oncogenic signaling. Nat Commun 16, 4833 (2025). 10.1038/s41467-025-60019-6

4 Godfrey, A. L. et al. JAK2V617F homozygosity arises commonly and recurrently in PV and ET, but PV is characterized by expansion of a dominant homozygous subclone. Blood 120, 2704–2707 (2012). 10.1182/blood-2012-05-431791

5 Grockowiak, E. et al. Different niches for stem cells carrying the same oncogenic driver affect pathogenesis and therapy response in myeloproliferative neoplasms. Nat Cancer 4, 1193–1209 (2023). 10.1038/s43018-023-00607-x

6 Li, R. et al. A proinflammatory stem cell niche drives myelofibrosis through a targetable galectin-1 axis. Sci Transl Med 16, eadj7552 (2024). 10.1126/scitranslmed.adj7552

7 Dunbar, A. J. et al. Jak2V617F Reversible Activation Shows Its Essential Requirement in Myeloproliferative Neoplasms. Cancer Discov 14, 737–751 (2024). 10.1158/2159-8290.CD-22-0952

8 Williams, N. et al. Life histories of myeloproliferative neoplasms inferred from phylogenies. Nature 602, 162–168 (2022). 10.1038/s41586-021-04312-6

9 Neubauer, H. et al. Jak2 deficiency defines an essential developmental checkpoint in definitive hematopoiesis. Cell 93, 397–409 (1998). 10.1016/s0092-8674(00)81168-x

10 Chen, M. et al. CRISPR/Cas12a-Based Ultrasensitive and Rapid Detection of JAK2 V617F Somatic Mutation in Myeloproliferative Neoplasms. Biosensors (Basel) 11 (2021). 10.3390/bios11080247

11 Baik, R., Wyman, S. K., Kabir, S. & Corn, J. E. Genome editing to model and reverse a prevalent mutation associated with myeloproliferative neoplasms. PLoS One 16, e0247858 (2021). 10.1371/journal.pone.0247858

12 Smith, C. et al. Efficient and allele-specific genome editing of disease loci in human iPSCs. Mol Ther 23, 570–577 (2015). 10.1038/mt.2014.226

13 Tsai, S. Q. et al. GUIDE-seq enables genome-wide profiling of off-target cleavage by CRISPR-Cas nucleases. Nat Biotechnol 33, 187–197 (2015). 10.1038/nbt.3117

14 Bozhilov, Y. et al. Reducing oxidative stress improves ex vivo polymer-based human haematopoietic stem and progenitor cell culture and gene editing. bioRxiv (2024). 10.1101/2024.09.17.613552

15 Khan, A. O. et al. Human Bone Marrow Organoids for Disease Modeling, Discovery, and Validation of Therapeutic Targets in Hematologic Malignancies. Cancer Discov 13, 364–385 (2023). 10.1158/2159-8290.CD-22-0199

16 Rodriguez-Meira, A. et al. Unravelling Intratumoral Heterogeneity through High-Sensitivity Single-Cell Mutational Analysis and Parallel RNA Sequencing. Mol Cell 73, 1292–1305 e1298 (2019). 10.1016/j.molcel.2019.01.009

17 Jakobsen, N. A. et al. Selective advantage of mutant stem cells in human clonal hematopoiesis is associated with attenuated response to inflammation and aging. Cell Stem Cell 31, 1127–1144 e1117 (2024). 10.1016/j.stem.2024.05.010

18 Zeng, A. G. X. et al. Single-cell Transcriptional Atlas of Human Hematopoiesis Reveals Genetic and Hierarchy-Based Determinants of Aberrant AML Differentiation. Blood Cancer Discov, OF1-OF18 (2025). 10.1158/2643-3230.BCD-24-0342

19 Young, I. C. et al. Differentiation of fetal hematopoietic stem cells requires ARID4B to restrict autocrine KITLG/KIT-Src signaling. Cell Rep 37, 110036 (2021). 10.1016/j.celrep.2021.110036

20 Li, Y. et al. Regulation of Alternative Polyadenylation Events by PABPC1 Affects Erythroid Progenitor Cell Expansion. bioRxiv (2025). 10.1101/2025.03.17.643825

21 Pagliarini, R., Shao, W. & Sellers, W. R. Oncogene addiction: pathways of therapeutic response, resistance, and road maps toward a cure. EMBO Rep 16, 280–296 (2015). 10.15252/embr.201439949

22 Frangoul, H. et al. Exagamglogene Autotemcel for Severe Sickle Cell Disease. N Engl J Med 390, 1649–1662 (2024). 10.1056/NEJMoa2309676

23 Olijnik, A. A. et al. Generating human bone marrow organoids for disease modeling and drug discovery. Nat Protoc 19, 2117–2146 (2024). 10.1038/s41596-024-00971-7

24 Schreurs, R., Baumdick, M. E., Drewniak, A. & Bunders, M. J. In vitro co-culture of human intestinal organoids and lamina propria-derived CD4(+) T cells. STAR Protoc 2, 100519 (2021). 10.1016/j.xpro.2021.100519

25 Clement, K. et al. CRISPResso2 provides accurate and rapid genome editing sequence analysis. Nat Biotechnol 37, 224–226 (2019). 10.1038/s41587-019-0032-3

26 Hao, Y. et al. Dictionary learning for integrative, multimodal and scalable single-cell analysis. Nat Biotechnol 42, 293–304 (2024). 10.1038/s41587-023-01767-y

