## Supplementary Figures for "Genome editing for treatment of *JAK2* V617F-driven myeloproliferative neoplasms"

**Supplementary Figure 1**

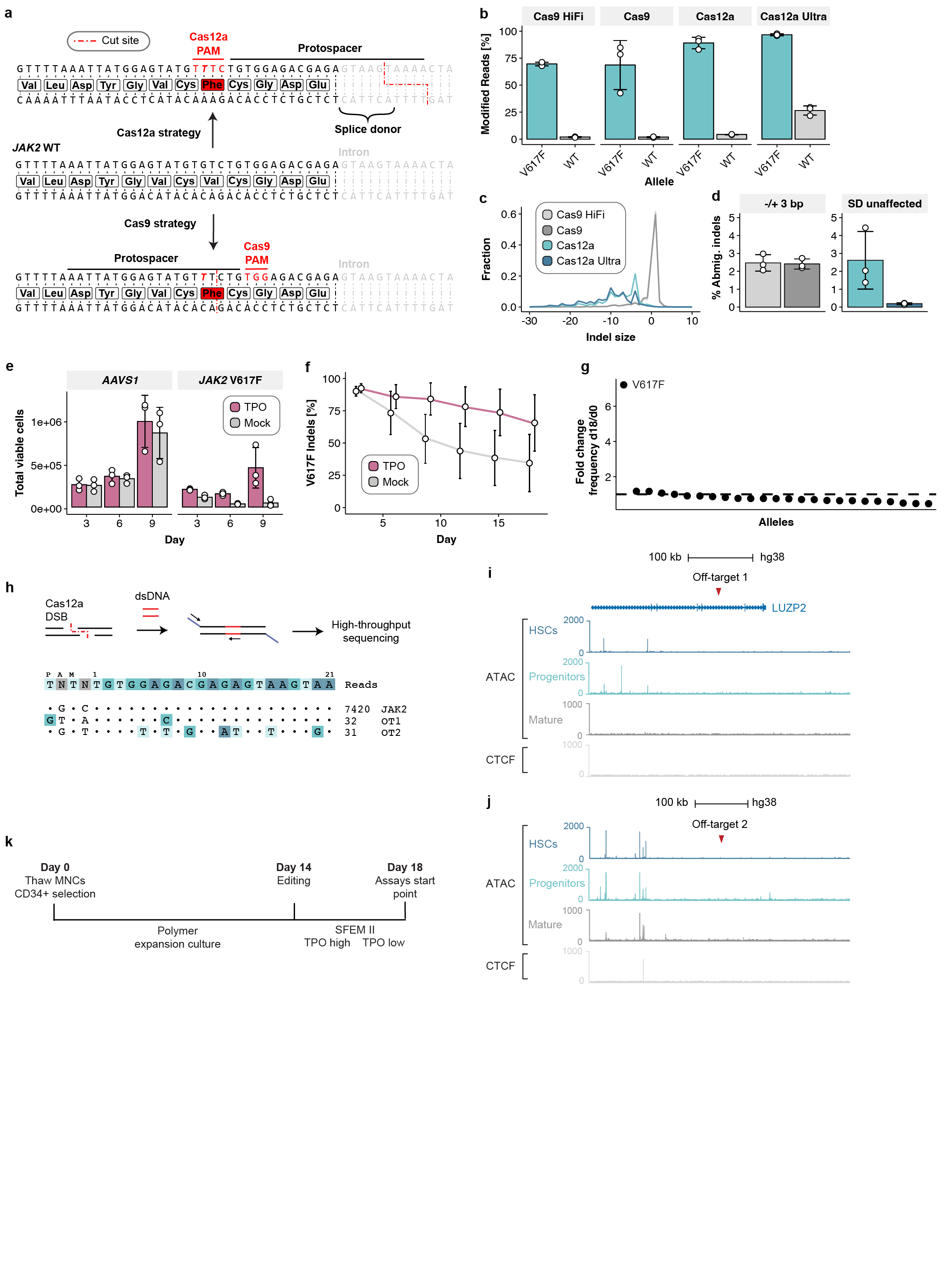
**Supplementary Fig. 1**. **a**, Schematic of the V617F locus and positioning of Cas9 and Cas12a allele-selective gRNAs. **b**, Editing efficiencies of WT and V617F alleles in SET-2 cells 72h after electroporation with Cas9 or Cas12a RNPs, determined via amplicon sequencing (n *=* 3 independent experiments, mean±s.d.). **c**, Indel size distribution with different nuclease (mean from n = 3 independent experiments). **d**, Percent reads with ambiguous indels from n = 3 independent editing experiments in SET-2 cells. For Cas9-based targeting, defined as reads with in-frame indels. For Cas12a-based targeting, defined as reads with indels that do not affect the splice donor (SD). **e**, Number of live SET-2 cells after editing for V617F or *AAVS1* control site, with or without supplementation of 20 ng/µL TPO (n *=* 6 independent experiments, mean±s.d.). **f**, Percent V617F reads with indels in SET-2 cells over time with or without supplementation of 20 ng/µL TPO (n *=* 3 independent experiments, mean±s.d.). **g**, Fold enrichment of editing products between day 18 and day 0 of culturing, with each dot representing a different allele that was present post editing (mean from n = 3 independent experiments). **h**, Schematic representation and GUIDE-Seq results for the Cas12a Ultra strategy in SET-2 cells. JAK2 is the on-target site and off-targets 1 and 2 (OT1/2) the identified off-targets. **i,j,** Genome browser views of OT1 and OT2 showing ATAC and CTCF data from various hematopoietic cell types from Corces et al. **k**, Schematic representation of culturing and editing protocol on patient-derived CD34+ cells.

**Supplementary Figure 2**

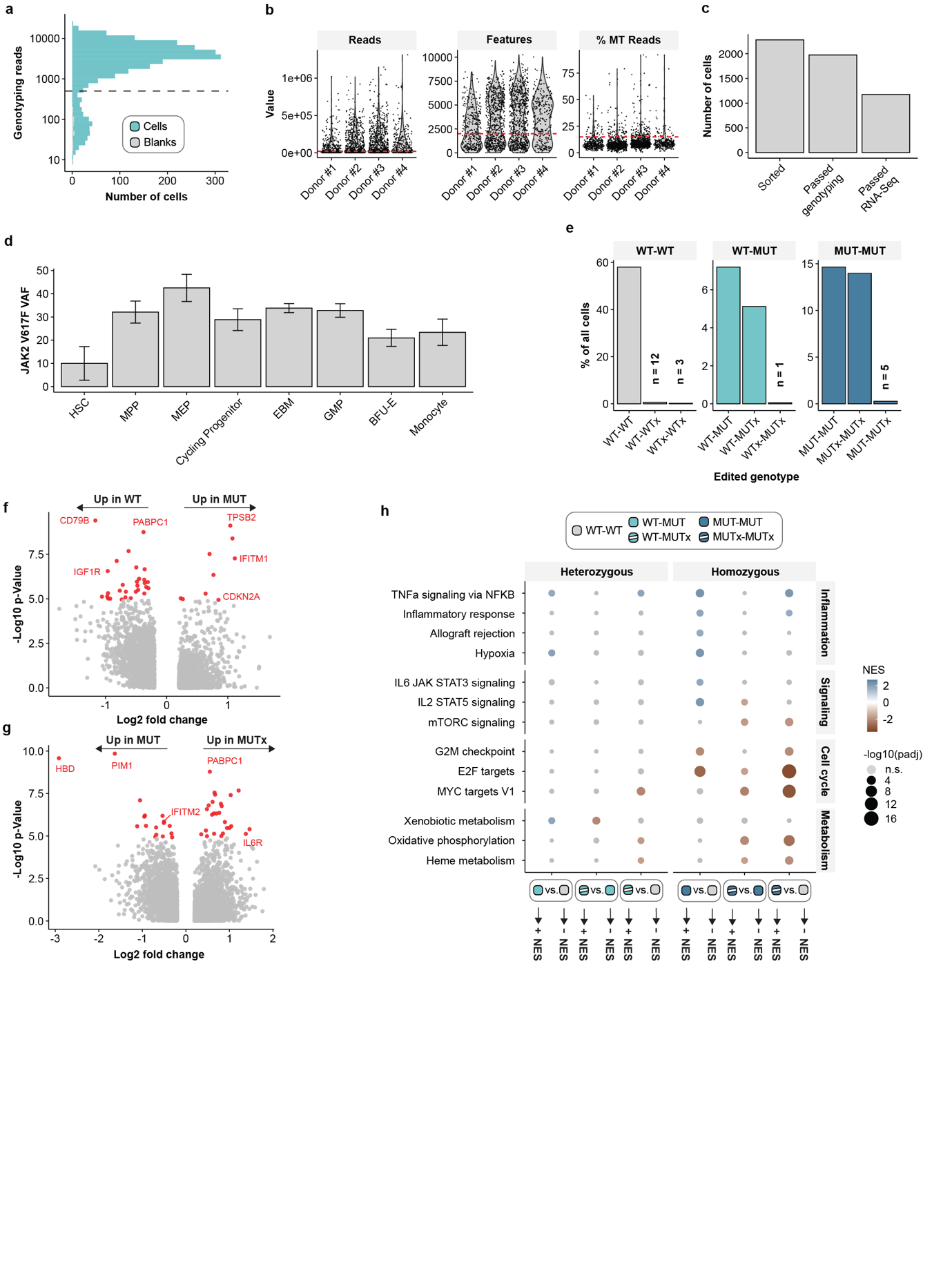

**Supplementary Fig. 2.** **a**, Numbers of reads per cell in the JAK2 genotyping library. Dashed line indicates filtering threshold. **b**, scRNA-Seq QC. Shown are number of reads, number of detected genes and percent reads mapping to mitochondrial (MT) genes for each cell in four donors. Red dashed lines indicate filtering threshold. **c**, Number of cells that passed the indicated QC steps. **d**, VAF by inferred cell type, calculated from genotyping results of the corresponding cells (mean±s.d.). **e**, Percent and number of cells with indicated genotype in the entire TARGET-Seq dataset. **f**, Volcano plot of differential expression between WT (i.e. all WT-WT) and MUT (i.e. both WT-MUT and MUT-MUT) cells. **g**, Volcano plot of differential expression between MUT (i.e. both WT-MUT and MUT-MUT) and MUTx (i.e. both WT-MUTx and MUTx-MUTx) cells. **h**, GSEA results split by heterozygous (i.e. WT-WT vs. WT-MUT vs. WT-MUTx) and homozygous (i.e. WT-WT vs. MUT-MUT vs. MUTx-MUTx) cells. All data obtained from a total of n = 4 independent patient samples.
